# Copy number variation and population-specific immune genes in the model vertebrate zebrafish

**DOI:** 10.1101/2023.08.23.554498

**Authors:** Yannick Schäfer, Katja Palitzsch, Maria Leptin, Andrew R. Whiteley, Thomas Wiehe, Jaanus Suurväli

## Abstract

Many species have hundreds of immune genes from the NLR family (Nucleotide-binding domain Leucine-rich Repeat containing). In plants they have a considerable amount of within-species variation, but not much is known about their variability in fishes. Here we captured and analysed the diversity of NLRs in zebrafish (*Danio rerio*) by sequencing 93 individuals from four wild and two laboratory strains. We found 1,560 unique NLR genes, and theoretical modelling revealed each wild population to have around 2,000. Only 100-550 were detected in each individual fish, and the observed variance of copy numbers differed among populations. Laboratory strains were found to have three times less NLRs than wild populations, and their genetic diversity was lower in general. Many NLRs showed no single nucleotide variation, but those that did showed evidence of purifying selection. Our study lays the groundwork for unraveling mechanisms driving the evolution of this large gene family in vertebrates.

**Significance statement:** We show here that the gene repertoires of vertebrates can be extremely variable, with different individuals having different genes. By sequencing one large family of immune receptors from 93 wild and laboratory zebrafish we found hundreds of novel gene copies, each only present in specific strains or specific individuals. Our observations can be explained by a combination of complex patterns of inheritance and a high rate of gene birth and death.

## Introduction

Every living being has a genetically encoded immune system, a highly plastic set of mechanisms used for defense against pathogens. Immune genes tend to evolve quickly and are often associated with a high degree of genetic variability. Many genes and proteins of the immune system are lineage-specific (limited to specific groups of animals, plants or other taxa), while others have defense roles in a wide range of species. In particular, proteins containing a large nucleotide-binding domain followed by smaller repeats have an immune function in animals, plants, fungi and bacteria alike (1–4). In land plants and animals, these repeats are often Leucine-rich Repeats (LRRs) and the proteins themselves are classified as NLRs (Nucleotide-binding domain Leucine-rich Repeat containing). Most genes conveying disease resistance in plants are NLRs, and different strains of thale cress *Arabidopsis thaliana* possess different repertoires of them (5). NLR repertoires can be also referred to as NLRomes, and a species-wide repertoire is called the ‘pan-NLRome’.

Animal NLRs (also known as NOD-Like Receptors) are only distantly related to those in plants (6, 7), but they are also essential for immunity. Some act as pathogen sensors or transcription factors (8), others are components or modulators of inflammasomes, large protein complexes that are assembled within cells as part of the response to biological or chemical danger (8). Most knowledge about animal NLRs comes from studies of humans and rodents, but their NLR repertoires (20-30 genes) are smaller than those of many other species (9–11). One NLR in mice (*Nlrp1*) has been shown to have different copy numbers in different strains, ranging from two to five (12).

In many fishes, studies have reported NLR repertoires in the range of 10-50 genes (13, 14). In others, hundreds of NLRs are present, including in the model species zebrafish (*Danio rerio*) (11, 15–18). The zebrafish reference genome contains nearly 400 NLR genes, two thirds of which are located on the putative sex chromosome (chromosome 4), in a genomic region associated with extensive haplotypic variation (19–22).

The majority of fish NLRs represent a fish-specific subtype that was originally labelled NLR-C (16), although they can be further divided into at least six groups based on structural and sequence similarities (18, 20). A schematic structure of proteins encoded by zebrafish NLR-C genes is presented on Figure 1A. All of them possess a FISNA domain (Fish-specific NACHT-associated domain), which precedes the nucleotide-binding domain NACHT and is encoded by the same large exon near the start of the protein (20). FISNA-NACHT is in some cases preceded by the effector domain PYD, but this is encoded by a separate exon (20). Additionally, many NLR-C proteins have a B30.2 domain (also known as PRY/SPRY) at the other end, separated from FISNA-NACHT by multiple introns and exons containing the LRRs (Figure 1A) (20). B30.2 is also found in other large immune gene families, such as the TRIM ubiquitin ligases (11, 20, 23).

**Figure 1.**
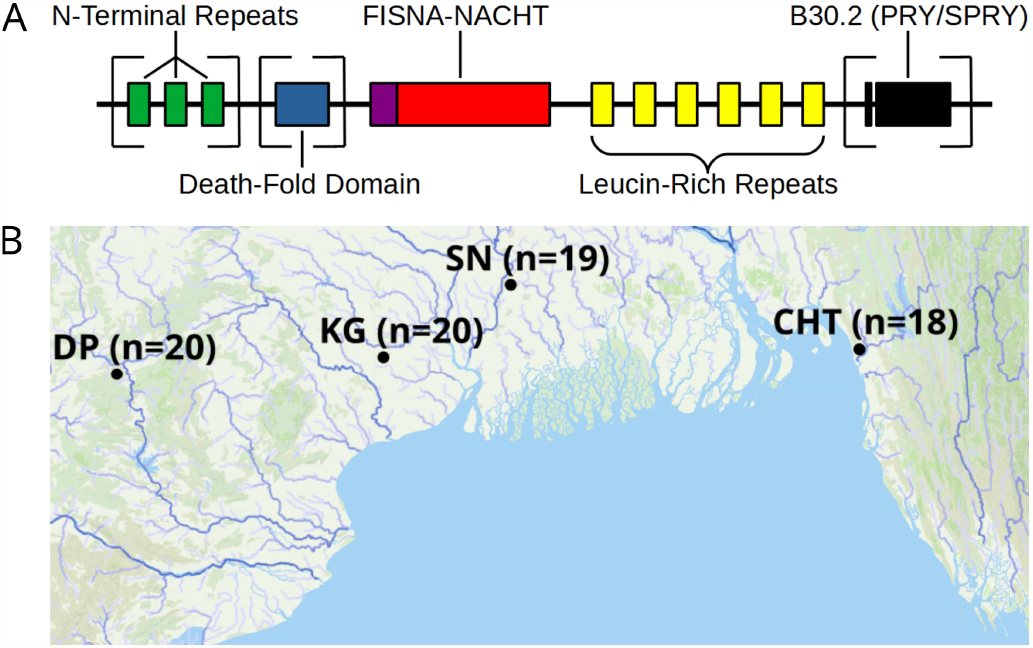
A: Generalized, schematic representation of the domain architecture of an NLR-C protein. Each box represents a translated exon. The N-terminal repeats, the death-fold domain as well as the B30.2 domain only occur in subsets of NLR-C genes. The number of N-terminal repeats and Leucine-Rich Repeats can vary. Domains that can be either present or absent in different NLRs are surrounded by square brackets. **B: Sampling sites for wild zebrafish**. All sites are located near the Bay of Bengal. Final sample sizes are indicated in parentheses. The map is based on geographic data collected and published by AQUASTAT from the Food and Agriculture Organization of the United Nations (27).

It is not known why fishes possess so many NLRs, how they evolve and how much within-species genetic variability they have. The previously observed repeated expansions and contractions of this family suggest it to have a high rate of gene birth and death (11). Studies have shown that viral and bacterial infections can induce the expression of specific fish NLRs (reviewed in (24)). Some have PYD or CARD domains and can even form inflammasomes similar to mammalian NLRs (25, 26). A species-wide inventory of major NLR exons (pan-NLRome) in a model species such as zebrafish would provide valuable insights into the evolution and diversity of this large immune gene family.

## Results

By adapting and modifying a protocol that combines bait-based exon capture with PacBio SMRT technology (28), we generated Circular Consensus Sequence (CCS) data for targeted parts of the immune repertoire from 93 zebrafish, representing four wild populations and two laboratory strains (Figure 1B, Supplementary Figure S1). With this approach, we aimed to sequence all exons in zebrafish that encode the nucleotide-binding FISNA-NACHT domains and all exons that encode B30.2 domains.

Our protocol used PCR with primers targeting ligated adapters to amplify the below-nanogram amounts of genomic DNA obtained from exon capture. This limited our fragment sizes to the lengths of what the polymerase was able to amplify. Zebrafish NLRs can have their exons spread out across tens of kilobases, so that we cannot know which exons belong to the same gene. However, we were able to use captured sequence surrounding the targeted exons to distinguish among near-identical coding sequences, and to separate NLR-associated B30.2 domains from B30.2 elsewhere in the genome (Supplementary Figure S3).

### The zebrafish pan-NLRome

We used an orthology clustering approach on NLR sequences assembled from all populations to create a reference set of NLRs (a pan-NLRome). This resulted in the identification of 1,560 unique sequences for FISNA-NACHT and 574 for NLR-associated B30.2 (NLR-B30.2). Additionally, we found 743 B30.2 domains not associated with NLRs. Nearly 10% of the sequences (150 FISNA-NACHT and 64 NLR-B30.2) contained pre-mature stop codons that were at least 10 amino acids from the end and lead to early truncation of the protein. 105 of the 1,560 FISNA-NACHT were preceded by an exon containing the N-terminal effector domain PYD. Nearly all of those (100 out of 105) were found in Group 1 NLR-C genes identified by the presence of the characteristic sequence motif GIAGVGKT (20).

Since the combination of FISNA and NACHT is only present in NLR-C, its count (1,560) can be considered equal to the total number of NLR-C genes in the data. We found each individual zebrafish to have at least one copy of 100-550 NLRs from the pan-NLRome (Figure 2). Additionally, each individual possesses at least one copy of 20-180 NLR-B30.2s (Supplementary Figure S4).

**Figure 2.**
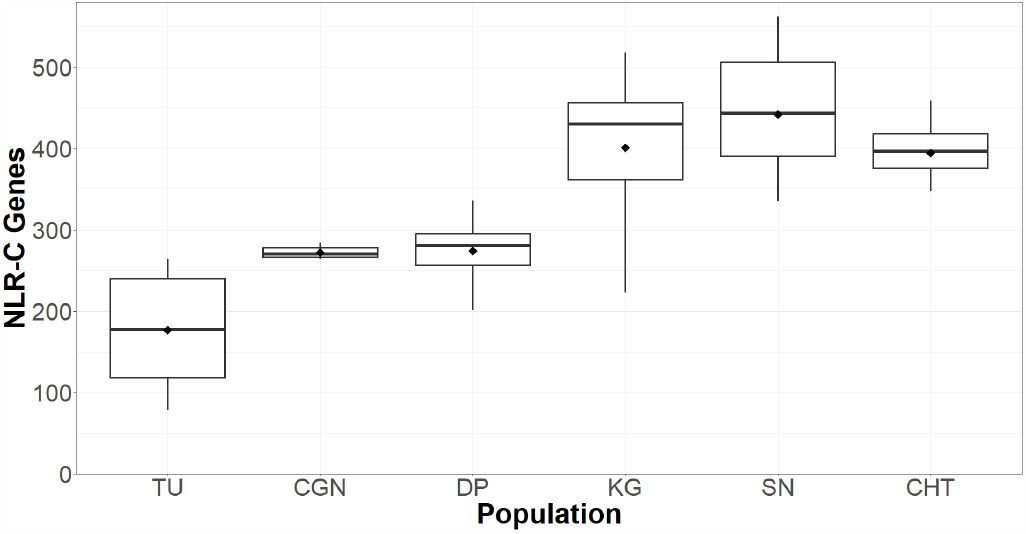
Total counts of NLRs found per individual. Black diamonds on the box plots denote means, horizontal lines denote medians.

### Copy number variation in the pan-NLRome

Aligning CCS reads to the pan-NLRome revealed a considerable amount of variability in the proportion of reads mapping to them, both between and within populations (Figure 3A). This can be interpreted as the gene being present in different copy numbers, Supplementary Figure S4). Furthermore, each NLR had its own distinct pattern of copy number variation, although generally the highest copy numbers were observed for wild populations KG, SN and CHT (Figure 3A). We also observed some sequencing batch-related differences, but the copy numbers differed even between individuals sequenced in the same batch.

**Figure 3.**
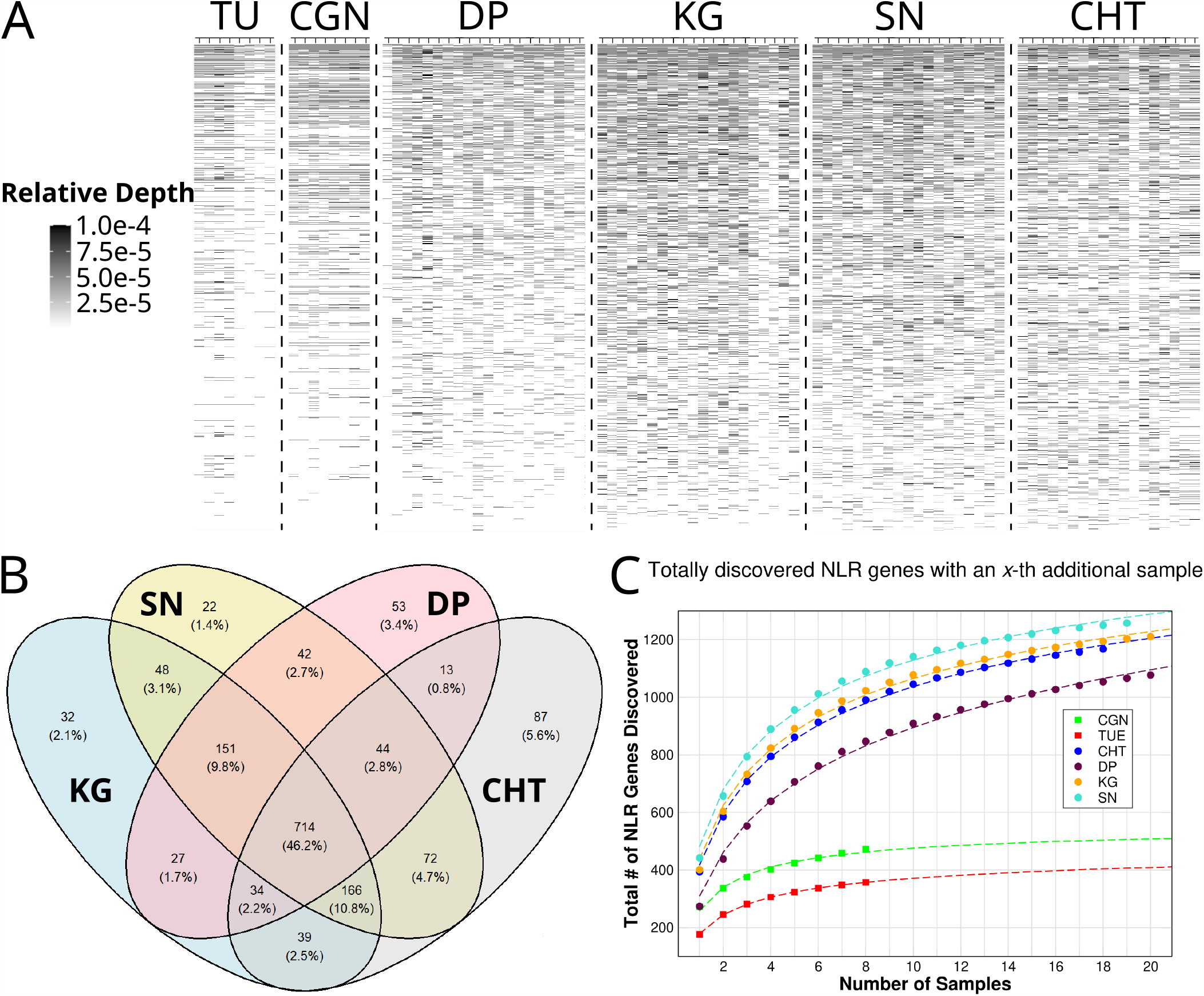
Copy number variation of NLR genes. A: Sequence data from each individual zebrafish (x-axis) was aligned to FISNA-NACHT exon sequences of the pan-NLRome (y-axis). Color intensity shows the proportion of data in a given fish that corresponds to each individual NLR. Darker colors can be interpreted as the exon potentially having multiple copies. Lighter colors indicate a single copy, white color means that the sequence was not found to be present. For clarity, only the 1,235 FISNA-NACHT for which at least one fish had 10 reads mapped to it are shown. B: Numbers of pan-NLRome sequences found in wild populations. Some are population-specific while others are shared by two or more populations. C: Estimated sizes of population-specific pan-NLRomes. Data points (filled circles and squares) show the average number of additionally discovered NLR genes (as identified via their FISNA-NACHT domain) when adding data from the *x*-th individual to a previously examined set of *x −* 1 individuals.

Of the 1,560 unique FISNA-NACHT and 574 NLR-B30.2 sequences, 714 FISNA-NACHT and 229 NLR-B30.2 (46% and 40%, respectively) were detected in at least one individual from all four wild populations (Figure 3B). These sequences can be thought of as the wild core set of NLRs. There were also NLR sequences shared between just two or three wild populations, and many were restricted to a single population (Figure 3B).

Fitting the number of NLRs discovered from each new individual to a harmonic function allowed us to estimate the sizes of the NLRomes of the populations (Figure 3C). We estimated a total of 520 and 570 NLRs in the laboratory strains TU and CGN, respectively. For the wild populations, we estimated four times as many, with 2,283 in KG, 1,896 in SN and 2,452 in CHT for KG, SN and CHT. For DP, the estimated pan-NLRome size was 25,284, but this is expected to be an artifact caused by low sequencing coverage for most samples of this population.

### Differences from the reference genome

NLRs sequenced in this study were often different from those present in the reference genome GRCz11. Even NLRs sequenced from the strain that the reference genome itself is based on (TU) did not always align well to it. When the exon itself did align, the intronic sequences surrounding it could often be very different from the reference. In numbers, only around 75% of NLRs occurring in TU fish aligned to the reference genome GRCz11 with high mapping qualities (Supplementary Figure S2). This number dropped even lower elsewhere - from 60-65% of NLRs in CGN which aligned well to the reference, down to only around 50% for the wild populations. The majority of NLRs that did not map well had a very poor mapping quality of 1 (Supplementary Figure S2). Moreover, there were 9 FISNA-NACHT and 10 NLR-associated B30.2 in the pan-NLRome which did not map anywhere in the reference genome.

### Purifying selection on single nucleotide variants

We used the pan-NLRome as a reference for identifying single nucleotide polymorphisms in the data. NLR sequence diversity was rare, with a large fraction of exons not having any variants in all populations. If variants were present, nucleotide diversity (*θ*_*π*_) was up to 0.016 and Watterson’s estimator (*θ*_*w*_) up to 0.021 (Figure 4A, B). In laboratory strains, genetic variability of FISNA-NACHT exceeded that of B30.2, but no such pattern was observed for wild populations. B30.2 exons of laboratory strains were also less variable than B30.2 from wild zebrafish (Figure 4B). The proportion of exons without any polymorphisms was much higher among FISNA-NACHT than among B30.2 (Figure 4C). The majority of variable NLR exons had *θ*_*π*_*/θ*_*w*_ ratios of less than 1 (Figure 4D), indicating an excess of rare alleles.

**Figure 4.**
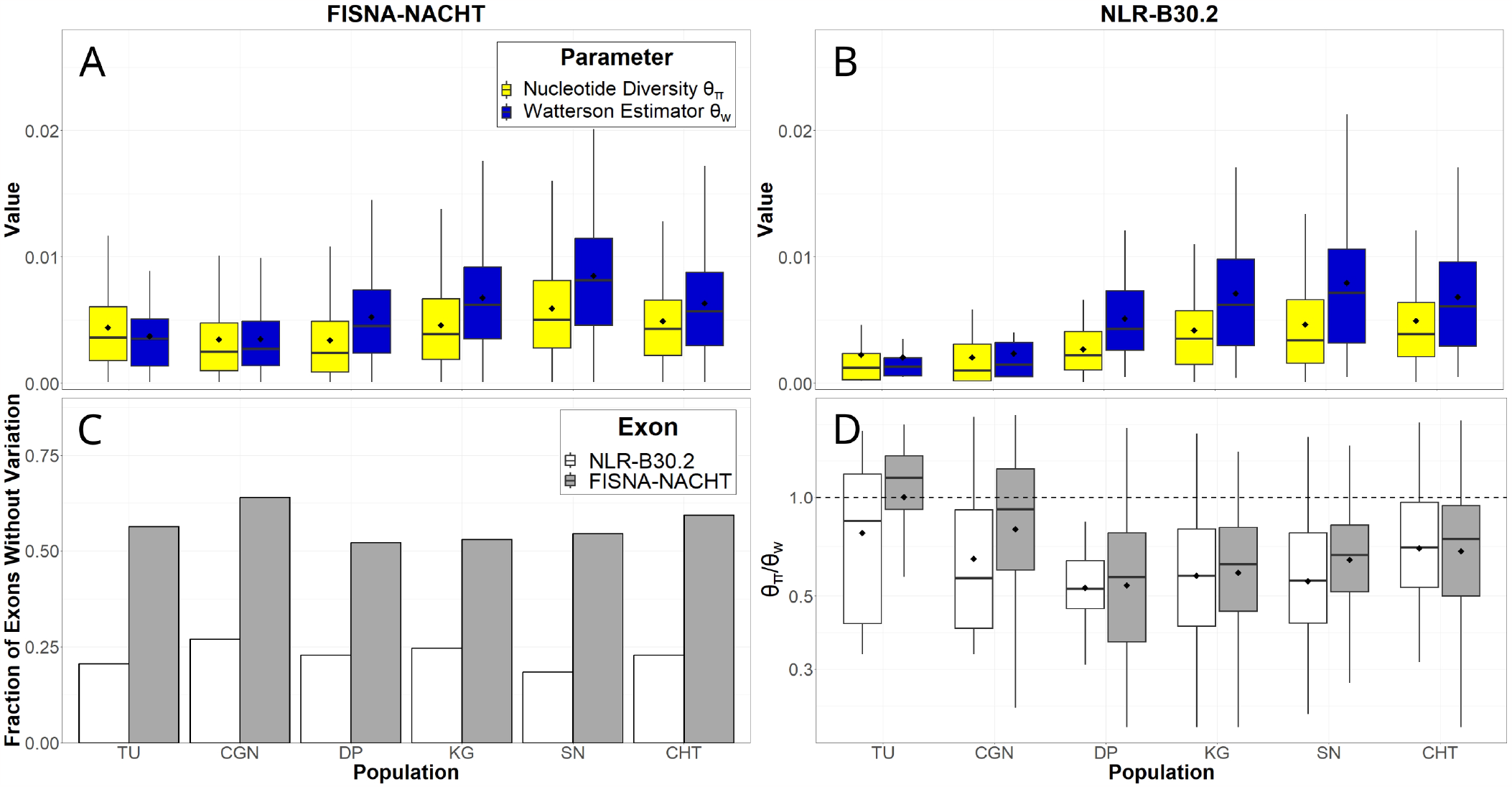
Single nucleotide variation of different NLR exons, shown by population. A: Nucleotide diversity (*θ*_*π*_) and Watterson estimator (*θ*_*w*_) for FISNA-NACHT exons. B: Nucleotide diversity (*θ*_*π*_) and Watterson estimator (*θ*_*w*_) for NLR-associated B30.2 exons. C: Proportion of exons without any single nucleotide variants. D: Ratio of *θ*_*π*_ */θ*_*w*_. Only exons with at least one variant are shown. The black, dotted line marks a ratio of 1. The black diamonds on box plots denote means, horizontal lines denote medians.

## Discussion

We sequenced and assembled the FISNA-NACHT and B30.2 exons of hundreds of NLRs from 93 zebrafish, thus capturing the diversity of this gene family in four wild populations and two laboratory strains. We found evidence that each genome from a wild individual contains only a fraction of more than 1,500 identified NLR copies. The number of unique NLRs per individual, each with one or more copies, ranged from around 100 to 550. While some of the lower numbers were likely underestimated due to low sequencing depths in specific samples and the population DP, the observed slow increase in newfound NLR gene copies per sample in all populations other than DP indicates that not many NLRs were missed.

Mathematical modelling based on a harmonic distribution suggested that the investigated laboratory strains possess roughly 500-600 NLRs while wild populations possess around 2,000. The pan-NLRomes are closed because sequencing more samples adds only few new NLRs to the total collection. However, to sequence all or almost all NLR genes, it would be necessary to sequence hundreds of individuals from each population. Besides, there are more, phylogeographically different populations - for example in Nepal and the Western Ghats (29) - which likely have tens if not hundreds of NLR copies not present in the populations we sequenced.

With a shift of focus towards pan-genomic studies (see (30–32)), future research on large multi-copy gene families should perhaps aim to generate gene databases similar to our pan-NLRome as starting points for deeper analyses. As shown for zebrafish NLRs, the availability of a single high quality reference genome may not be sufficient to estimate the true diversity of large gene families.

### Properties of the zebrafish NLRome

We have previously demonstrated a substantial reduction in single nucleotide variation in zebrafish laboratory strains compared to wild populations (33). Here we showed that the copy numbers of the NLRome are similarly reduced. Similarly, we also find laboratory strains to have less presence/absence variation than wild populations. The initial bottleneck and the reduced amount of pathogenic challenges in a laboratory environment could lead to a rapid loss of expendable genes, especially if unequal recombination is frequent (34).

Our predicted number of NLRs in a population is on the same scale as the 2,127 NLRs found in the NLRome of *Arabidopsis thaliana* (5). Structural similarities of NLRs in plants and animals are thought to be the result of convergent evolution (7), but it appears that the similarities extend beyond domain structure. While the zebrafish and *A. thaliana* face different environments and pathogenic challenges, their NLRs appear to have undergone analogous evolutionary trajectories, such as the formation of isolated gene clusters (35). Copy numbers vary greatly between *A. thaliana* accessions (35), which is something we also observe in zebrafish. The sizes of NLR repertoires differ between zebrafish individuals in at least three of the wild populations. Whether the low number of NLR genes in the fourth wild population, DP, is an artifact caused purely by low coverage or also by evolutionary factors remains unclear at this time. In *A. thaliana*, 464 conserved, high confidence orthogroups were previously identified, 106-143 of which were defined as the core NLRome because they were found in a large subset of accessions (5). In wild zebrafish, we found a core NLRome consisting of 714 NLR genes, each of which was detected in at least one individual from all wild populations. Only half of those (52.5%, 375 in total) could be confidently mapped to the reference genome. We postulate that as immune genes, many of the core NLR genes are likely conserved between populations because they provide a fitness advantage in the defense against common pathogens. However, some of them may be retained due to neutral processes such as genetic drift. The additional NLRs shared among only some of the wild populations, and the population-specific NLRs may represent local adaptations to ecological niches. Additionally, there could be functional redundancy within the NLRome, so that different individuals have different NLRs with the same functional role. This idea is further supported by the fact that most wild fish had around 400 NLRs and only a subset of the core NLRome.

Although we mainly used the counts of FISNA-NACHT orthogroups to estimate total numbers of NLRs, we also analyzed the B30.2 exons of NLR-C genes. In general, NLR-associated B30.2 exons exhibit patterns of copy number variation that are similar to those seen for FISNA-NACHT. For example, about half of the B30.2 sequences are found in all wild populations, similar to the “core” set of FISNA-NACHT exon sequences.

### Function of NLR-C genes in zebrafish

The fact that hundreds of NLR gene copies are maintained in zebrafish, together with hints of purifying selection, suggests that the evolution of these genes is driven by significant selective pressures. Although the expression of fish NLRs is often induced by pathogen exposure (reviewed in (24)), the exact function of most zebrafish NLR-C genes is unclear. Zebrafish has inflammasomes that make use of specific NLRs ((26, 36, 37), (25)), but the majority of NLR-C genes do not possess the necessary N-terminal effector domains (CARD or PYD) (38).

One possibility is that the zebrafish NLR-C genes simply act as intracellular pathogen sensors. Accordingly, the LRRs that all NLRs contain could be used to bind specific pathogen-associated molecular patterns. Group 4 NLR-C genes for example ((20)) contain no clear domains other than FISNA-NACHT and the LRRs themselves. However, in many cases the LRRs are followed by an additional domain, B30.2, for which the exact role in NLRs is unknown. It is possible that it takes up the role of the effector in at least some of the cases, or even has a direct role in pathogen recognition. In other proteins, it has been reported to interact with immune repertoire components such as *β*2-microglobulin, interferon-regulatory factors or caspases (39–41). In any case, the consistent presence of B30.2 exons in NLR-C genes as well as their excess of rare alleles point to a selective benefit of the B30.2 exon.

### What drives the copy number differences?

There are at least two mechanistic explanations for the extensive copy number variation seen among zebrafish populations: first, it could be attributed to a high degree of haplotypic variation. Large DNA fragments contain different sets of genes and gene copies, similar to the zebrafish MHC loci (42). Extensive haplotypic variation occurs on the long arm of chromosome 4, the location containing over two thirds of all NLRs in zebrafish (21). Such segregating haplotype blocks would explain the existence of the core NLRome, but not the frequent presence of genes that occur only in a single individual.

Alternatively, the evolution of NLR-C genes could be primarily driven by duplication events (43) and secondarily by gene conversion (16). Gene duplications are caused by unequal recombination, transposon activity or whole genome/chromosome duplications (44, 45). The arrangement of NLR-B30.2 genes in clusters on the long arm of chromosome 4 suggests that tandem duplication via unequal crossing-over (34) played the most important role in the expansion. Since there are many transposable elements on the long arm of chromosome 4 (19), it would be reasonable to assume that at least some of them have assisted in the local expansion and transfer of NLR exons and genes to chromosomes other than chromosome 4.

It is tempting to speculate that chromosome 4 could be a source of NLRs which continuously generates new copies. However, to maintain a stable genome size, gene gains must be balanced by gene loss. NLR-C genes may be lost via accumulation of random mutations due to a lack of selective pressure and loss-of-function mutations, but they may also be lost through unequal recombination. This mechanism would allow only NLR genes contributing to the functionality of the immune system to be kept, while others would disappear.

Although the NLRs of plants and animals arose independently (7), their evolutionary paths appear to have much in common, making them a spectacular example of convergent evolution. In plants, tandem duplication is thought to be the primary driver of NLR gene expansion (43), but they are also often associated with transposable elements. If the diversity of unrelated NLR genes in such distantly related species is driven by common molecular mechanisms, then the same mechanisms might also act on NLRs of other phylogenetic clades and even on unrelated large gene families, such as odorant receptors (46).

### Conclusion

This study advances our understanding of the evolutionary dynamics affecting very large gene families. The sheer amount of copy number variation that appears to be present in a single gene family of zebrafish is staggering, with different individuals each having numerous genes that are not present in all others. This can only be caused by diversity-generating mechanisms that are active even now. In this study, we have laid the groundwork for future studies investigating the molecular basis and evolutionary mechanisms contributing to the diversity of large, vertebrate gene families.

### Code and data availability

NLR reads are available in the NCBI Sequence Read Archive (BioProject PRJNA966920). Scripts are available on GitHub (https://github.com/YSchaefer/pacbio_zebrafish).

## Materials and methods

### Samples

Wild zebrafish from four sites in India and Bangladesh (Figure 1B, Supplementary Table S1) had been collected in the frame of other projects (29, 47). Laboratory zebrafish from the Tübingen (TU) and Cologne (CGN) strains were provided by Dr. Cornelia Stein (University of Cologne). All samples were stored in 95% ethanol until use. Tail fins from 20 fish per wild population and 8 fish per laboratory strain were used as starting material for the subsequent steps.

### DNA extraction, exon capture and sequencing

Genomic DNA was extracted with kits from QIAGEN (MagAttract HMW kit) and MACHEREY-NAGEL (Nucleospin Tissue Kit), followed by shearing with red miniTUBEs on the Covaris M220 ultrasonicator. Nicks in the DNA were repaired with PreCR Repair Mix (New England Biolabs). Samples were barcoded with the NEBNext Ultra II DNA Library Prep Kit, then pooled together in batches of 4 or 8 (details provided in Supplementary Methods). RNA baits for the exon capture (Daicel Arbor Biosciences) were custom-designed to target immune genes of interest (mainly NLRs). Exon capture and PacBio library preparation were both done according to a protocol adapted from Witek and colleagues (28). Libraries were sequenced at the Max Planck Genome Centre, with PacBio Sequel and Sequel II. Additional details are provided in Supplementary Methods, the pooling and sequencing scheme is presented in Supplementary Table S2. PCR protocols and the sequences of qPCR primers used to confirm enrichment success can be found in Supplementary Tables S3, S4 and S5.

### Read processing, mapping and clustering

Raw sequences were demultiplexed with `lima`. Consensus sequences of DNA fragments with at least three passes (CCS reads) were inferred with `ccs`, followed by PCR duplicate removal with `pbmarkdup`. All read mapping was done with `pbmm2` (v.1.3.0), a PacBio wrapper for `minimap2` (48). `lima`, `ccs`, `pbmarkdup` and `pbmm2` were all provided by Pacific Biosciences. Mapped files were processed and filtered with `samtools` (v1.7) (49). *De novo* assemblies were generated with `Hifiasm` (v0.15.4-r347) (50). Tools from the `HMMER` suite (v3.2.1) (51) were used to detect the presence of NLR-associated sequences. Contigs containing FISNA-NACHT or B30.2 were sorted into orthoclusters using `get_homologues` (build x86 64-20220516) (52) and `blastn` (v2.11.0+) (53). Orthoclusters for which `pbmm2` did not align any CCS reads to the representative sequence with at least 95% identity were excluded from further analyses. Further details are provided in Supplementary Methods.

### Modelling

To estimate the full size of each wild population’s NLR repertoire, we modelled the increase in the total number of identified NLR exon sequences when adding observations from a new individual from a population to a set of already analysed individuals (as shown in (54)). Briefly, given a sample of *n* individuals, there are

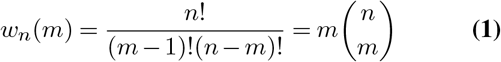

ways to choose *m* − 1 individuals from the entire sample and add another - not yet chosen - one. For each *m* we calculated the average increase in the number of identified exon sequences over all possible choices of individuals. Based on these numbers we fitted the non-linear function

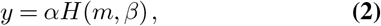

where *H*(*m, β*) is the generalized harmonic number with parameter *β*, i.e.

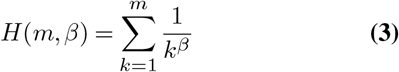

in which *β* = 1 is equivalent to the expected number of segregating sites in a standard coalescent (Equation 4). Here, we may reinterpret lineage splits as duplication events and a parameter *β >* 1 reflecting an increase of the duplication rate towards the present. Importantly, if *β >* 1, the series in equation (3) converges, and its limit may be interpreted as the size of a closed NLRome. The NLRome is open if *β* ≤ 1. Values of the fitted parameters and saturation limits are presented in Supplementary Table S6.

### Genetic diversity

Single nucleotide variants in each fish were identified from the .bam output of `pbmm2` by using `deepvariant` (r1.0) (55) with the PacBio model. Joint genotyping of the individual samples was done with `glnexus` (v1.2.7-0-g0e74fc4) (56) with its deepvariant-specific setting. Per-site *θ*_*π*_ of the NLR exons was calculated with `vcftools` (v0.1.16) (57). *θ*_*w*_ was calculated via the formula

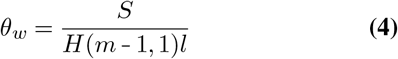

where *S* is the number of segregating sites and *l* is the length of the exon (1,761 for FISNACHT and 540 for B30.2).

### Data visualization

Plots, heat maps and Venn diagrams were created in RStudio (v2022.07.2) with R (v4.2.1) using `ggplot2` (v3.3.6) and `ggvenn` (v0.1.9) or `xmgrace` (v5.1.25; https://plasma-gate.weizmann.ac.il/Grace/). Final processing of the images was done in Inkscape (v1.1.2; https://inkscape.org/).

## Conflict of Interest statement

The authors declare no conflict of interest.

## Author contributions

JS co-designed and coordinated the study, prepared sequence libraries from laboratory zebrafish, contributed to computational analyses and co-wrote the manuscript. YS did the majority of computational analyses and co-wrote the manuscript. KP extracted DNA and prepared sequence libraries from wild zebrafish. ML co-designed the study. TW co-designed the study and did the copy number modelling. ARW obtained wild zebrafish samples. All authors contributed to the final manuscript with their commentaries and feedback.

## Acknowledgements

The authors are thankful to Emilia Martins and Anuradha Bhat for their contributions to wild sample collection, and Cornelia Stein and the lab of Mathias Hammerschmidt for laboratory zebrafish. The authors are grateful to Bruno Hüttel, who provided advice and support during library construction and sequenced all samples at the Max Planck Genome Centre. We also acknowledge the contributions of Philipp Schiffer (help with writin
g the initial project proposal), Lisa Vogelsang (assistance with lab work), Robert Fürst and Anna Rottmann (management of computational infrastructure). This work was funded by a grant to TW and ML in the frame of the priority program SPP1819 of the German Research Council (DFG). JS was additionally funded by a National Sciences and Engineering Council of Canada (NSERC) Discovery Grant to Colin Garroway.

## Supplementary Information

**Figure S1.**
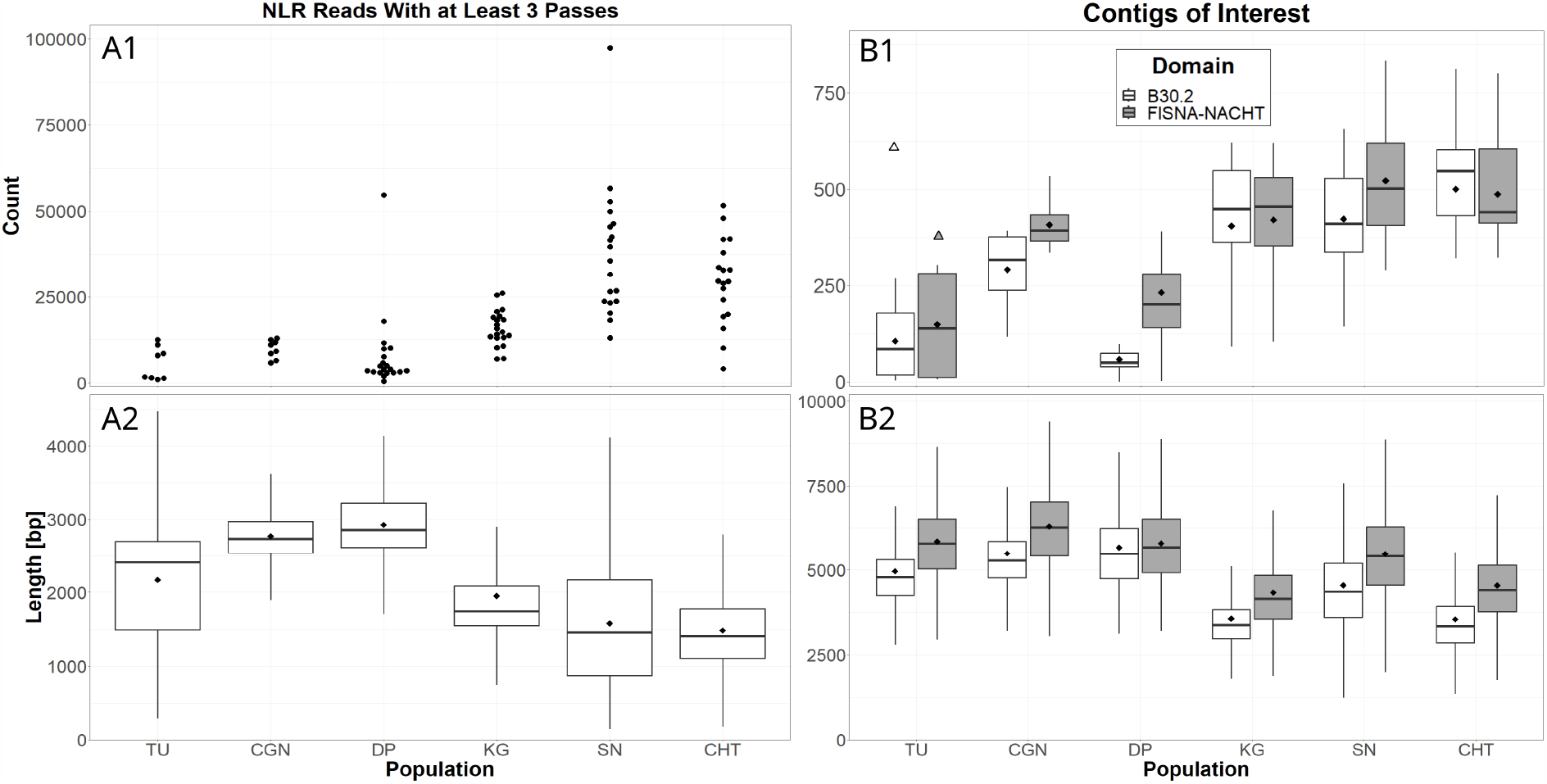
Sequencing and assembly statistics of CCS reads from NLR exons. A1: Absolute numbers of CCS reads from NLR exons per sequenced individual. A2: Lengths of the CCS reads that map to NLR genes. B1: Absolute numbers of assembled contigs containing an NLR exon per sequenced individual. Triangles above TU mark the numbers of NLR exons found in the reference genome. B2: Lengths of the individual assembled contigs that contain an NLR exon. Outliers not shown in the boxplots. The black diamonds on boxplots denote means, horizontal black lines denote medians.

**Figure S2.**
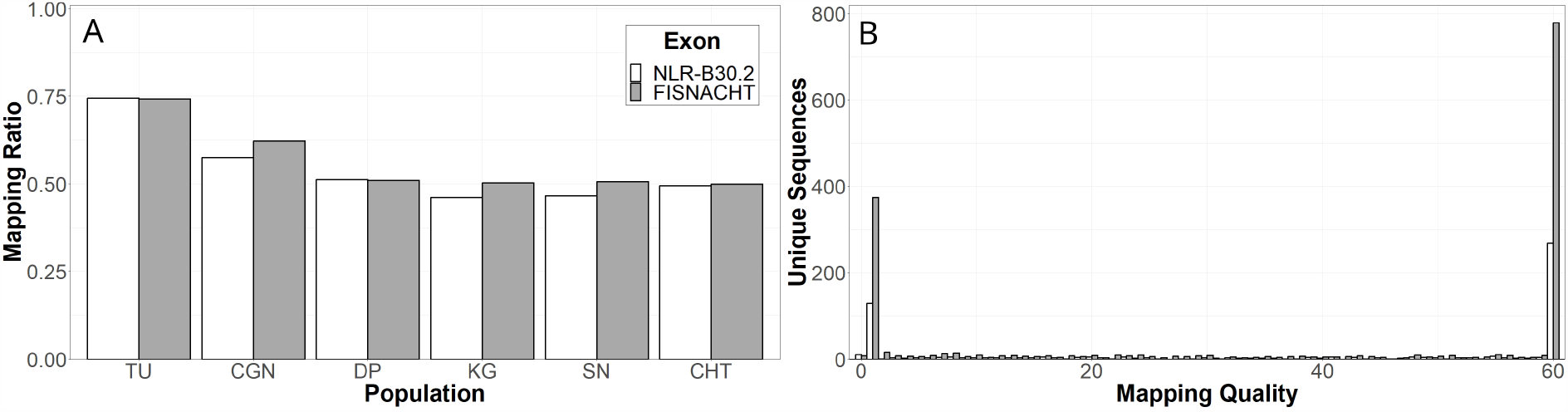
NLRome vs reference genome GRCz11. A: Proportions of unique FISNACHT and NLR-B30.2 sequences that were successfully mapped to the reference genome GRCz11 with a mapping quality of 60, by population. B: Distribution of mapping qualities for all unique NLR sequences that aligned to GRCz11, showing that most map either with very high (60) or very low quality.

**Figure S3.**
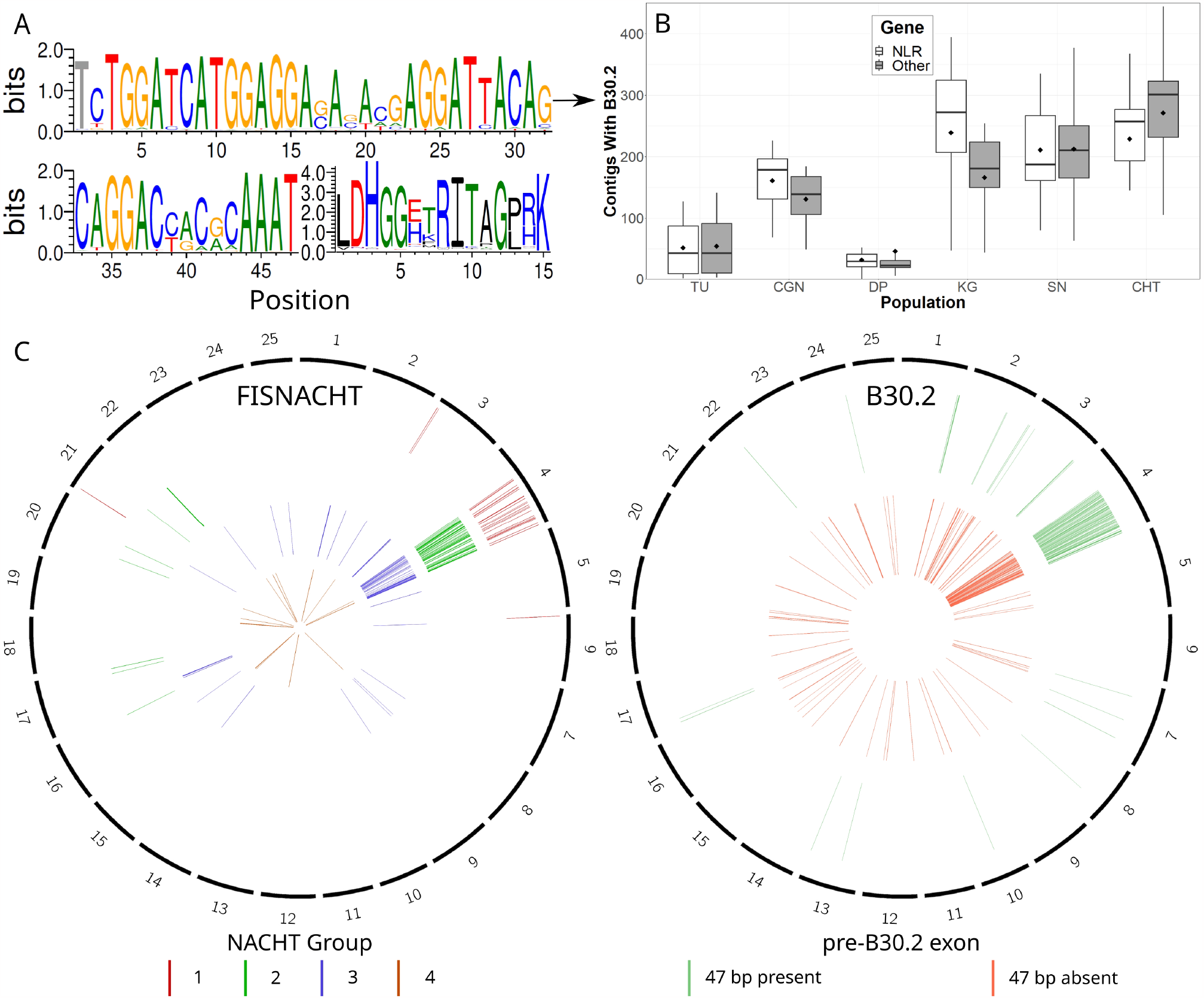
Small highly conserved exon that precedes B30.2 in zebrafish NLRs. A: Nucleotide sequence logo (on top, continued on bottom left) and amino acid logo (on bottom right) of the NLR-specific exon. The first nucleotide of the exon was removed to generate the correct amino acid translation. Logos were created with Weblogo, the height of each base represents its information content in bits (58). B: Absolute numbers of contigs containing a B30.2 exon per sequenced individual, split by presence/absence of the NLR-specific exon. The black diamonds on boxplots mark the means. C: Circos plots of the genomic distribution of FISNACHT sequences (left) by NACHT group and B30.2 sequences (right) by presence of the NLR signature motif.

**Figure S4.**
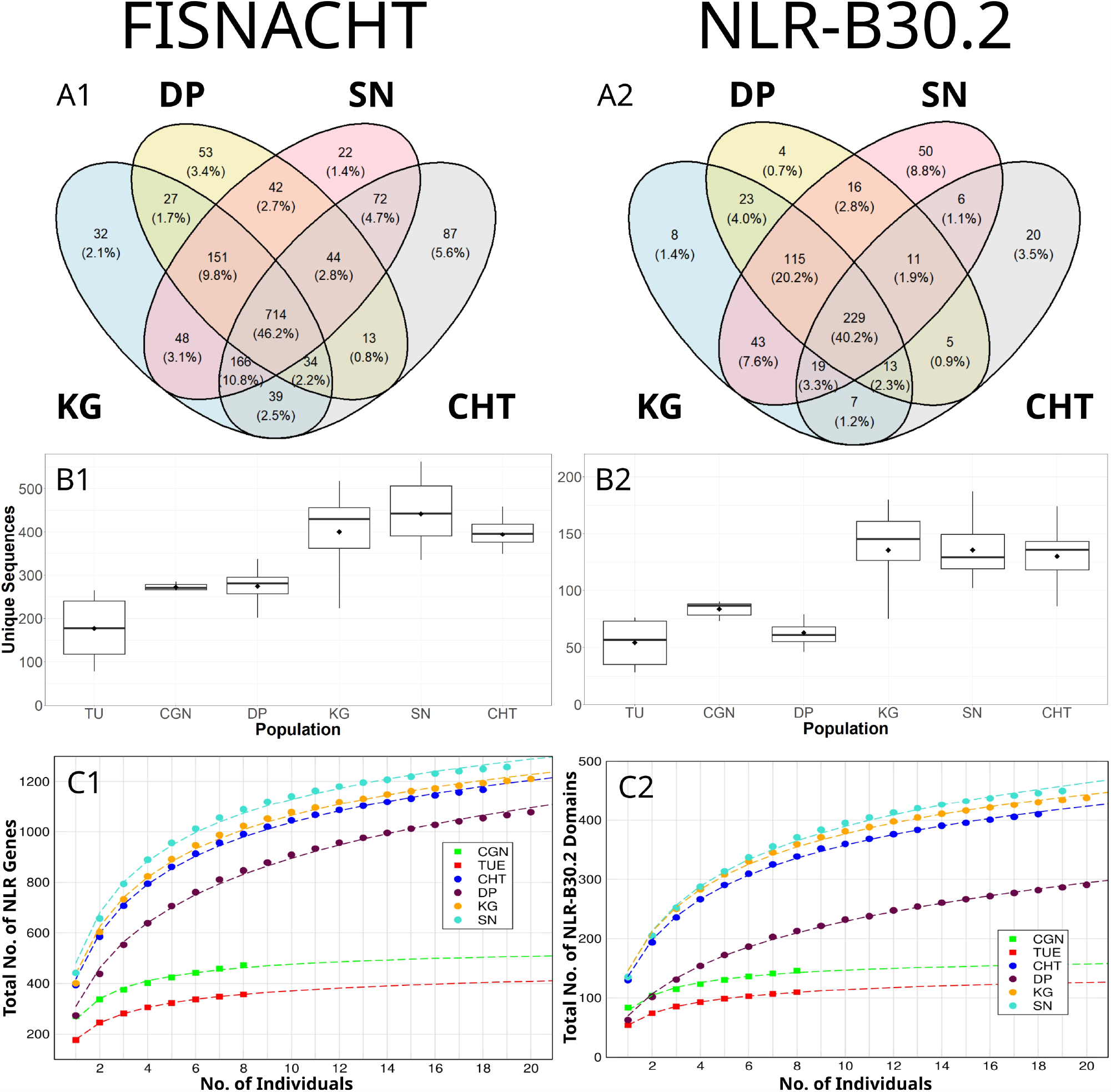
Copy number variation of NLR genes. A1, A2: Numbers of private and shared NLR sequences in wild populations. B1, B2: Numbers of unique NLR sequences (each with one or more copies per individual) found in fish of the sequenced strains. Black diamonds on the box plots denote means, horizontal lines denote medians. C1, C2: Population-specific pan-NLRomes and sets of NLR-B30.2 domains. Data points (filled circles and squares) show the average number of additionally discovered NLR genes (as identified via their FISNACHT domain) when adding data from the *x*-th individual to a previously examined set of *x −* 1 individuals. The dotted lines represent the result of non-linear curve fitting to the function 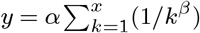, the generalized *x*-th harmonic number (detailed in Materials and Methods).

**Table S1.** Zebrafish samples sequenced in this study. Although initially there were 20 samples from each wild population, some did not yield enough material for succesful sequencing.

| Name | Origin | Coordinates | Samples | Sequenced |
| --- | --- | --- | --- | --- |
| Tübingen (TU) | N/A (pet shop) | N/A | 8 | 8 |
| Cologne (CGN) | N/A (pet shop) | N/A | 8 | 8 |
| Dandiapalli (DP) | Odisha, India | N22.22155<br>E84.79430 | 20 | 20 |
| Leturakhal (KG) | West Bengal, India | N22.26189<br>E87.27881 | 20 | 20 |
| Santoshpur (SN) | West Bengal, India | N22.93765<br>E88.55311 | 20 | 19 |
| Chittagong (CHT) | Bangladesh | N22.47400<br>E91.78300 | 20 | 18 |

**Table S2.** Sequencing scheme for the zebrafish samples. Libraries sequenced after the introduction of an improved (Long Run) sequencing chemistry are marked with LR. Samples that yielded no data after sequencing are marked with asterisks.

| Individuals | Library | Sequencing Technology |
| --- | --- | --- |
| TU01, TU02, TU03, TU06 | TU L1 | PacBio Sequel<br>(SMRT Cell 1M v2) |
| TU08, TU10, TU12, TU14 | TU L2 | PacBio Sequel<br>(SMRT Cell 1M v2) |
| CGN1, CGN2, CGN3, CGN4 | CGN L1 | PacBio Sequel<br>(SMRT Cell 1M v2) |
| CGN5, CGN6, CGN7, CGN8 | CGN L2 | PacBio Sequel<br>(SMRT Cell 1M v2) |
| DP07, DP09, DP10, DP12 | DP L1 | PacBio Sequel<br>(SMRT Cell 1M v2) |
| DP15, DP20, DP23, DP24, DP25, DP28, DP31, DP34 | DP L2 | PacBio Sequel<br>(SMRT Cell 1M v2, LR) |
| DP03, DP05, DP13, DP16, DP21, DP29, DP31, DP33 | DP L3 | PacBio Sequel<br>(SMRT Cell 1M v2, LR) |
| KG35, KG41, KG42, KG43 | KG L1 | PacBio Sequel<br>(SMRT Cell 1M v2) |
| KG03, KG05, KG07, KG12, KG14, KG15, KG18, KG19 | KG L2 | PacBio Sequel<br>(SMRT Cell 1M v2, LR) |
| KG20, KG22, KG24, KG26, KG29, KG32, KG33, KG44 | KG L3 | PacBio Sequel<br>(SMRT Cell 1M v2, LR) |
| SN21, SN23, (SN24*), SN26 | SN L1 | PacBio Sequel<br>(SMRT Cell 1M v2) |
| SN03, SN04, SN08, SN09, SN10, SN11, SN12, SN24 | SN L2 | PacBio Sequel II<br>(SMRT Cell 8M, LR) |
| SN13, SN14, SN15, SN16, SN17, SN18, SN19, SN20 | SN L3 | PacBio Sequel II<br>(SMRT Cell 8M, LR) |
| CHT19, CHT23, CHT26, CHT28 | CHT L1 | PacBio Sequel<br>(SMRT Cell 1M v2) |
| CHT01 - CHT07, (CHT13*) | CHT L2 | PacBio Sequel II<br>(SMRT Cell 8M, LR) |
| CHT08, CHT10 - CHT12, CHT14 - CHT16, (SN25*) | CHT L3 | PacBio Sequel II<br>(SMRT Cell 8M, LR) |

**Table S3.**
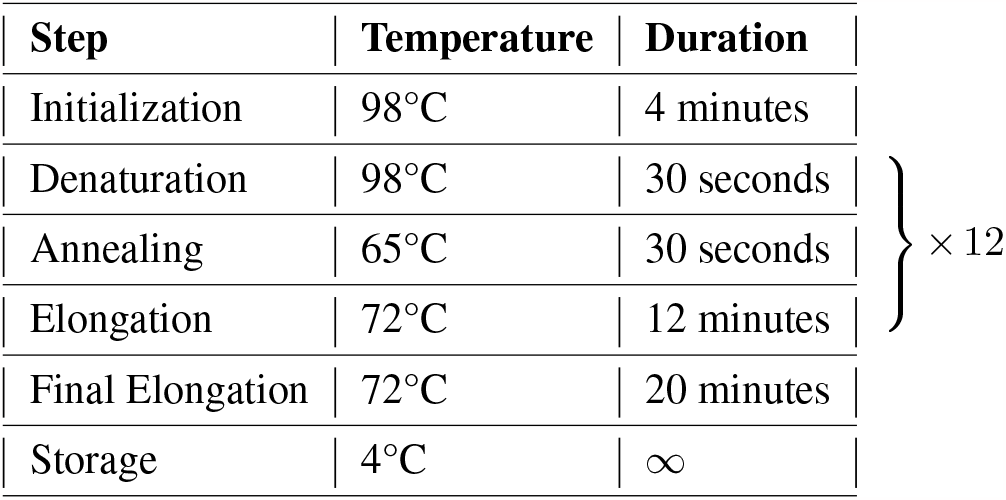
PCR program used for barcoding. For library amplification, the same program was used with 26 or 31 cycles

**Table S4.** qPCR program for the evaluation of enrichment efficiency.

| Step | Temperature | Duration |
| --- | --- | --- |
| Initialization | 95°C | 12 minutes |
| Denaturation | 95°C | 15 seconds |
| Annealing | 65°C | 20 seconds |
| Elongation | 72°C | 20 minutes |

**Table S5.** Sequences of qPCR primers used for evaluation of target enrichment.

| Gene | Direction | Sequence |
| --- | --- | --- |
| il1 | + | 5'-tgg-tga-acg-tca-tca-tcg-cc-3' |
| il1 | - | 5'-tcc-agc-acc-tct-ttt-tct-cca-a-3' |
| foxo6 intron | + | 5'-agt-tct-gtg-tgg-gaa-cag-gg-3' |
| foxo6 intron | - | 5'-gtg-cat-ctt-tag-cgt-tgg-ct-3' |
| NLR group 1 | + | 5'-cct-gac-aca-ggt-caa-caa-aac-a-3' |
| NLR group 1 | - | 5'-gat-tgt-ctt-ttc-ctt-cag-ccc-ag-3' |
| NLR group 2 | + | 5'-tgg-att-ggg-ctg-aag-gga-aa-3' |
| NLR group 2 | - | 5'-agg-ttc-agt-cct-tta-gtc-tct-gg-3' |
| NLR group 3 | + | 5'-ctg-ctg-gag-gtg-aaa-gat-cag-ac-3' |
| NLR group 3 | - | 5'-gat-tgt-tga-gca-gtg-agc-agg-a-3' |
| NLR group 4 | + | 5'-tac-ctg-gac-aag-aca-aag-cca-3' |
| NLR group 4 | - | 5'-ctc-ctt-ctc-ttc-agc-cca-gtc-3' |

**Table S6.** Values of fitted parameters and saturation limits for FISNACHT and NLR-B30.2 exons, by population.

| Population | FISNACHT |  |  | NLR-B30.2 |  |  |
| --- | --- | --- | --- | --- | --- | --- |
| | $\alpha$ | $\beta$ | Limit | $\alpha$ | $\beta$ | Limit |
| TU | 178.274 | 1.43356 | 519.548 | 53.8579 | 1.40774 | 164.73 |
| CGN | 257.207 | 1.62786 | 569.367 | 78.7156 | 1.61283 | 177.246 |
| DP | 309.14 | 1.01231 | 25284 | 69.3609 | 0.87454 | ∞ |
| KG | 436.761 | 1.2152 | 2288.41 | 145.715 | 1.1418 | 1113.23 |
| SN | 479.892 | 1.26093 | 2152.12 | 145.548 | 1.10183 | 1514.35 |
| CHT | 416.712 | 1.18893 | 2451.81 | 135.677 | 1.11911 | 1218.54 |

## Supplementary methods

### DNA extraction

High Molecular Weight (HMW) DNA from laboratory zebrafish was extracted from caudal fin clips using the QIAGEN MagAttract HMW DNA extraction kit. HMW DNA from wild zebrafish was extracted from caudal fin clips using the NucleoSpin Tissue Kit from MACHEREY-NAGEL with the following adjustments. Tissues other than muscle were removed before DNA extraction with forceps. The incubation time of the Proteinase K treatment was changed from 1-3 hours to 10-15 minutes. An RNAse A treatment step was included by incubating with 400 µg RNAse A (Sigma-Aldrich) for 2 minutes at room temperature. All DNA samples were quantified and quality checked with Qubit 3.0 (ThermoFisher Scientific), 0.8% agarose gels and the 4200 TapeStation Electrophoresis System (Agilent Technologies). DNA extraction failed for one of the 20 CHT samples, but was succesful for the other 95 fin clips.

### Shearing and barcoding

High molecular weight (HMW) DNA was sheared into 1.5 to 6 kb fragments with the red miniTUBEs of the Covaris M220 ultrasonicator. Quality control after shearing was performed using the 4200 TapeStation Electrophoresis System (Agilent Technologies). The obtained DNA fragments were size selected with 0.4x AMPure XP beads (Beckmann Coulter Inc) to exclude fragments smaller than 1.5 kb. For wild zebrafish samples, a DNA damage repair step was included in order to repair any possible DNA damage resulting from long periods of storage (particularly important for the older CHT samples). The repair step was carried out with PreCR Repair Mix (New England Biolabs).

DNA fragments were barcoded with the NEBNext Ultra II DNA Library Prep Kit (New England Biolabs) and NEBNext Multiplex Oligos for Illumina, Index Primers Set 1 (New England Biolabs). The manufacturer’s standard protocol was followed until the amplification step for the enrichment of barcode-ligated fragments. At this stage, the recommended amplification protocol (PCR program) was modified to suit large DNA fragments (Table S3) and the high fidelity Kapa polymerase (Kapa HiFi Hotstart Readymix, Kapa Biosystems) was used. The resulting barcoded DNA was purified and size-selected two more times, first with 0.5x AMPure XP beads and then with 0.4x AMPure XP beads. The amount of DNA was quantified with Qubit and quality checked with gel electrophoresis on a 0.8% agarose gel. The samples were then pooled with each pooled sample containing barcoded DNA of either 4 fish (CGN, TU, first library of each wild population) or 8 fish (the remaining libraries of the wild populations).

### NLR capture with hybridization baits

Target enrichment was carried out using the MYbaits v3 customized target enrichment kit for Next Generation Sequencing (MYcroarray, now part of Daicel Arbor Biosciences). The bait set contained nearly 20,000 unique 120-bp biotinylated RNA molecules in equimolar amounts. Most of the baits were designed to specifically bind to the FISNACHT and B30.2 exons in the genome, but we also targeted other genes of interest. Bait hybridization and target enrichment for each pooled sample was performed according to the MYbaits manual version 3.02, with half the amount of baits and reagents used for the 4-fish pools than for the 8-fish pools. Following an overnight incubation of the pooled DNA samples with RNA baits, bait-bound DNA fragments were extracted from the solution with DB MyOne Streptavidin C1 beads (ThermoFisher Scientific). The enriched libraries were subsequently amplified with P5 and P7 primers (Illumina) by running 26 cycles of the program described in Table S4. If the DNA yield was less than 1,000 ng afterwards (measured by Qubit), five more PCR cycles were added. Enrichment success was evaluated by qPCR, using 5x HOT FIREPol EvaGreen qPCR Mix Plus (ROX) (Solis BioDyne) and primers specific for the FISNACHT exons from each of the four groups of NLRs (Table S4, Table S5). The gene IL-1 was used as a single copy control. All primers were custom-ordered from biomers.net GmbH. The qPCR experiment was deemed succesful if a strong enrichment could be seen for all NLR groups, weaker enrichment for il-1, and no enrichment for the random intron. After this, the sample was selected for subsequent PacBio library construction and purified with 0.7x Ampure PB Beads (Pacific BioSciences).

### Library construction and sequencing

The final libraries were prepared with the SMRTbell Template Prep Kit 1.0-SPv3 (Pacific BioSciences). At the ligation step, the recommended amount of PacBio adapters was increased from 1 to 5 µl per 40 µl total reaction volume and the reaction was incubated over night at room temperature. For the SN and CHT libraries in pools of eight (see Table S2), barcoded PacBio adapters were used instead of regular ones. The product codes for barcodes were BC1001 and BC1002 for CHT, BC 1003 and BC1004 for SN.

The first libraries (TU, CGN, 4 DP and 4 KG samples) were size selected to 2-8 kb with the BluePippin pulsed field electrophoresis system (Sage Science). The following libraries were size selected to 1.5-8kb.

All sequencing was done at the Max Planck Genome Centre, located at the Max Planck Institute for Plant Breeding in Cologne. All TU, CGN, DP and KG zebrafish, as well as 4 CHT and 4 SN samples were sequenced with 1M v2 SMRT Cells of the Sequel instrument (Pacific BioSciences). The rest of the samples (all with barcoded adapters) were multiplexed together and sequenced with an 8M SMRT Cell of the much higher throughput Sequel 2 instrument (Pacific BioSciences). One of the already sequenced SN samples (SN24) was also resequenced in this run, as it yielded no data in the first one. Furthermore, Pacific BioSciences upgraded their kits with a superior polymerase after we had sequenced TU, CGN and the first 4 samples of each wild population; all samples other than those were sequenced with their LR (Long Run) polymerase.

### Read processing and assembly

Raw data were de-multiplexed and stripped of primer/adapter sequences with `lima` from Pacific Biosciences. For the samples sequenced with the PacBio Sequel I, the parameters `-enforce-first-barcode-split-bam-named -W 100` were used with `lima` v1.0.0 for the runs without the LR polymerase. For Sequel runs with the LR polymerase, lima v1.8.0 and v1.9.0 were used with the same parameters. To remove PacBio barcodes from the data produced on Sequel II, `lima` v1.11.0 was used with parameters `-split-bam-named -peek-guess` and for the subsequent removal of NEBNext barcodes, the parameters were changed to `-enforce-first-barcode-split-bam-named -peek-guess`. Consensus sequences of all DNA fragments with a minimum of three passes (henceforth referred to as CCS reads) were calculated using `ccs` (v4.2.0, Pacific Biosciences) with default parameters. PCR duplicates were identified and flagged with `pbmarkdup` (v1.0.0, Pacific Biosciences) with default parameters, then excluded from downstream analyses. Any chimeric reads containing a primer sequence in the middle were identified with `blastn` (v2.11.0+) (53) and removed. The filtered CCS reads were assembled into contigs for each fish separately using `hifiasm` (v0.15.4-r347) (50) with default parameters.

### NLR identification

In addition to the known NLR genes extracted from Ensembl (Release 107), the reference genome was translated in all frames using `transeq` (from EMBOSS:6.6.0.0) (59) and searched for further NLRs using `hmmsearch` from hmmer (v3.2.1) (51), without bias correction and with the Hidden Markov Model (HMM) profiles for zf_FISNA-NACHT and zf_B30.2 from (18). Each position in which the zf_FISNACHT model found a hit with a maximum i-Evalue of 1*e* − 200 and a minimum alignment length of 500 aa was considered a FISNACHT exon. The filtering thresholds for B30.2 exons were an i-Evalue of 1*e* − 5 and a minimum alignment length of 150 aa.

To distinguish CCS reads of NLR genes from other CCS reads, the CCS reads of each fish were mapped against the reference genome GRCz11 using `pbmm2` (v1.3.0) with preset ccs. CCS reads which mapped within a known NLR gene or one found with our HMM-based approach were considered NLR reads.

Contigs containing NLR exon sequences were identified by translating all contigs of each fish in all frames with `transeq` (from EMBOSS:6.6.0.0) and subsequently searching for FISNACHT and B30.2 domains using `hmmsearch` from hmmer (v3.2.1) without bias correction and the Hidden Markov Models (HMMs) zf_FISNA-NACHT and zf_B30.2 again. The HMM-based approach was chosen for the contigs in particular because we assumed that there would be NLR sequences in the data which are absent in the reference genome and therefore might not be mapped. The approach enabled us to include all FISNACHT and B30.2 exon data found by (18) in our searches, optimizing the search sensitivity.

By examining all NLRs annotated in the reference genome, we found a highly conserved 47 bp exon preceding B30.2 to be present in most NLR-B30.2 genes (NLRs containing a B30.2 domain), but not in other NLRs nor in most other B30.2-containing genes. B30.2 exons from NLRs were distinguished from B30.2 elsewhere in the genome by generating a Hidden Markov Model for the 47-bp exon based on the blast hits and searching the contigs for matches to this model with `hmmsearch` from hmmer (v3.2.1). The model was created with `hmmbuild` from hmmer (v3.2.1).

The FISNA-NACHT and B30.2 orthoclusters were postprocessed after `get_homologues` as follows: whenever an orthocluster contained more than one contig, a consensus sequence for the cluster was created from all those contigs with `cons` from EMBOSS:6.6.0.0. These consensus sequences and the contig sequences of the singleton clusters made up the representative sequences of the orthoclusters. Some representative sequences were reversed with `revseq` from EMBOSS:6.6.0.0 so that all exons were in the same orientation. The representative sequences were then blasted against each other using `blastn` (v2.12.0+) with default parameters and output format 6. In cases in which 98% and at least 3 kb of a representative sequence matched another with at least 98% identity, the two clusters they represented were fused into a new cluster by combining their contigs and generating a new consensus sequence from them. This process was conducted twice and reduced the number of FISNA-NACHT clusters from the initial 4,743 to 2,008 and B30.2 clusters from 14,879 to 2,635.

The bam files produced by mapping the NLR reads of each fish separately to the representative sequences of the orthoclusters were filtered using `samtools` (v1.7) (49). If the representative sequence had at least one primary alignment (SAM flag 0 or 16) with length > 1 kb, mapping quality 60 and no more than 9 soft-clipping bases at both ends of the mapped read, the orthocluster was assumed to occur in the respective fish.

Circular genome plots were created with `circos` (v 0.69-8) (60) running on Perl 5.036000.

## Notes

### Competing Interest Statement

The authors have declared no competing interest.

### Summary of Updates

The paper was rewritten to be more concise, without changing the message of it. Repetitive figures and long detailed descriptions in Methods were moved to Supplementary Data and replaced with shorter versions in the main text. In addition, a significance statement was added after the abstract. The formatting of the file was also changed, and now uses the Overleaf / LaTeX template zHenriquesLab-StyleBioRxiv, with added line numbers. Supplementary data can now be found at the end of the file, after references.

